# Bayesian Inference with Incomplete Knowledge Explains Perceptual Confidence and its Deviations from Accuracy

**DOI:** 10.1101/2020.09.18.304220

**Authors:** Koosha Khalvati, Roozbeh Kiani, Rajesh P. N. Rao

**Affiliations:** Paul G. Allen School of Computer Science and Engineering, University of Washington, Seattle, WA 98115, USA; Center for Neural Science, New York University, New York, NY 10003, USA; Department of Psychology, New York University, New York, NY 10003, USA; Neuroscience Institute, NYU Langone Medical Center, New York, NY 10016, USA; Center for Neurotechnology, University of Washington, Seattle, USA

## Abstract

In perceptual decisions, subjects infer hidden states of the environment based on noisy sensory information. Here we show that both choice and its associated confidence are explained by a Bayesian framework based on partially observable Markov decision processes (POMDPs). We test our model on monkeys performing a direction-discrimination task with post-decision wagering, demonstrating that the model explains objective accuracy and predicts subjective confidence. Further, we show that the model replicates well-known discrepancies of confidence and accuracy, including the hard-easy effect, opposing effects of stimulus volatility on confidence and accuracy, dependence of confidence ratings on simultaneous or sequential reports of choice and confidence, apparent difference between choice and confidence sensitivity, and seemingly disproportionate influence of choice-congruent evidence on confidence. These effects may not be signatures of sub-optimal inference or discrepant computational processes for choice and confidence. Rather, they arise in Bayesian inference with incomplete knowledge of the environment.

## 1 Introduction

Making decisions about hidden states of the environment based on noisy sensory information is critical for survival. Should an animal continue to graze after hearing a rustling sound? Was the sound due to a stalking predator or the wind? The outcome of such perceptual decisions is both a choice and an expectation of success known as confidence. Confidence plays a key role in guiding behavior in complex environments (Dayan and Daw, 2008; Kiani and Shadlen, 2009; Purcell and Kiani, 2016a; Vickers, 1979) and is often critical for modeling behavior and understanding its neural mechanisms in such environments (Beck et al., 2008; Drugowitsch et al., 2014; Fetsch et al., 2012; Kanitscheider et al., 2015; Purcell and Kiani, 2016b; Seilheimer et al., 2014). However, unlike sensory choices and their accuracy that are usually easy to measure, confidence is a subjective quality difficult to measure reliably, unless special experimental procedures are employed (Kepecs et al., 2008; Kiani and Shadlen, 2009; Kiani et al., 2014; Middlebrooks and Sommer, 2012; Persaud et al., 2007; Peters et al., 2017; Zylberberg et al., 2016; Fleming et al., 2018). Experiments that make such measurements have often revealed systematic discrepancies between subjective confidence reports and experimentally measured accuracy (Pouget et al., 2016; Juslin et al., 1997; Zylberberg et al., 2012; Lak et al., 2014; Zylberberg et al., 2016; Peters et al., 2017; Odegaard et al., 2018). These discrepancies have been occasionally interpreted as evidence for suboptimality of the decision-making process or for disparate processes for computing choice and confidence. Contrary to those interpretations, we show that a Bayesian framework with optimal inference but incomplete knowledge about the environment can explain choice accuracy, confidence, and their discrepancies in experimental measurements.

Our model extends Partially Observable Markov Decision Processes (POMDPs) (Kaelbling et al., 1998), which assume that subjects optimize a reward function by adjusting their beliefs about stimulus identity and the best choice based on two factors: sensory observations and prior knowledge about environmental states (Rao, 2010; Huang et al., 2012; Huang and Rao, 2013), which are learned from past experience. The model enables us to simulate temporal update of belief for a sequence of sensory observations. These belief updates generate explicit links between the decision maker’s confidence and choice accuracy. We demonstrate the precision of our predictions about choice confidence by testing them on monkeys performing a direction discrimination task with post-decision wagering (Kiani and Shadlen, 2009), where both choice accuracy and confidence were measured.

In addition to explicitly linking confidence and accuracy, our model explains well-known discrepancies between these two measurements. Some discrepancies arise in an optimal decision-making process when the decision maker has incomplete knowledge about the environment and needs to resolve uncertainties about the reliability of observations. Others seem to exist from an experimenters’ perspective because the exact information used by the subject is hidden to the experimenter. Our POMDP model explains commonly observed discrepancies between accuracy and confidence such as the “hard-easy” effect (Juslin et al., 1997; Drugowitsch et al., 2014; Sanders et al., 2016), higher confidence with increased variability of sensory observations despite reduction of accuracy (Fetsch et al., 2014; Zylberberg et al., 2016), different confidence ratings in simultaneous versus sequential reports of choice and confidence (Kiani et al., 2014; Sanders et al., 2016; van Den Berg et al., 2016), discrepancy between sensitivities of accuracy and confidence (*d′* vs. meta-*d′*) (Maniscalco and Lau, 2012; Fleming and Daw, 2017), and the seemingly larger effect of choice-congruent observations on confidence reports (Zylberberg et al., 2012; Peters et al., 2017).

We conclude by showing that the Bayesian inference component of our POMDP model can be implemented by the neural mechanisms that integrate evidence toward a decision bound, consistent with drift diffusion models (DDMs) (Ratcliff et al., 2016) or more generally, models based on bounded-accumulation of evidence. The POMDP model commits to a choice when the value of the expected improvement of accuracy with new observations is less than the cost of making those observations. We show that this termination criterion uniquely maps to a time-varying decision bound for integration of evidence in the DDM (see also Huang and Rao (2013)). Such time-varying bounds match past behavioral studies (Ratcliff and Rouder, 1998; Reddi et al., 2003) and can be implemented by the urgency signals observed in electrophysiological recordings (Churchland et al., 2008; Purcell and Kiani, 2016b; Cisek et al., 2009). Overall, the neural implementation of inference and choice in our POMDP framework is both simple and plausible.

## 2 Results

We developed and tested our model using behavioral data from monkeys performing a direction discrimination task with post-decision wagering (Fig. 1a) (Kiani and Shadlen, 2009). On each trial, monkeys observed a patch of randomly moving dots (Britten et al., 1992) and decided about the net direction of motion. The difficulty of the decision was varied randomly from trial to trial by changing the percentage of coherently moving dots (the “motion strength” or “coherence”) and the duration of the motion stimulus (Fig. 1b). The stimulus was followed by a delay period and at the end of the delay, the fixation point disappeared (Go cue), signaling the monkey to report its choice with a saccadic eye movement. On a random half of trials, the monkey was given only the right and left direction targets. Choosing the correct motion direction (right target for rightward motion and left target for leftward motion) resulted in a large reward (a large drop of juice) but choosing the incorrect target resulted in no reward and a short timeout. On the other half of trials, the monkey was offered a third target, in addition to the direction targets, in the middle of the delay period. This third target was a sure-bet option. The monkey could choose either the direction targets or the sure-bet after the Go cue. Choosing the sure-bet target guaranteed reward but the magnitude of the reward (volume of the juice) was smaller than that for choosing the correct direction target.

**Figure 1:**
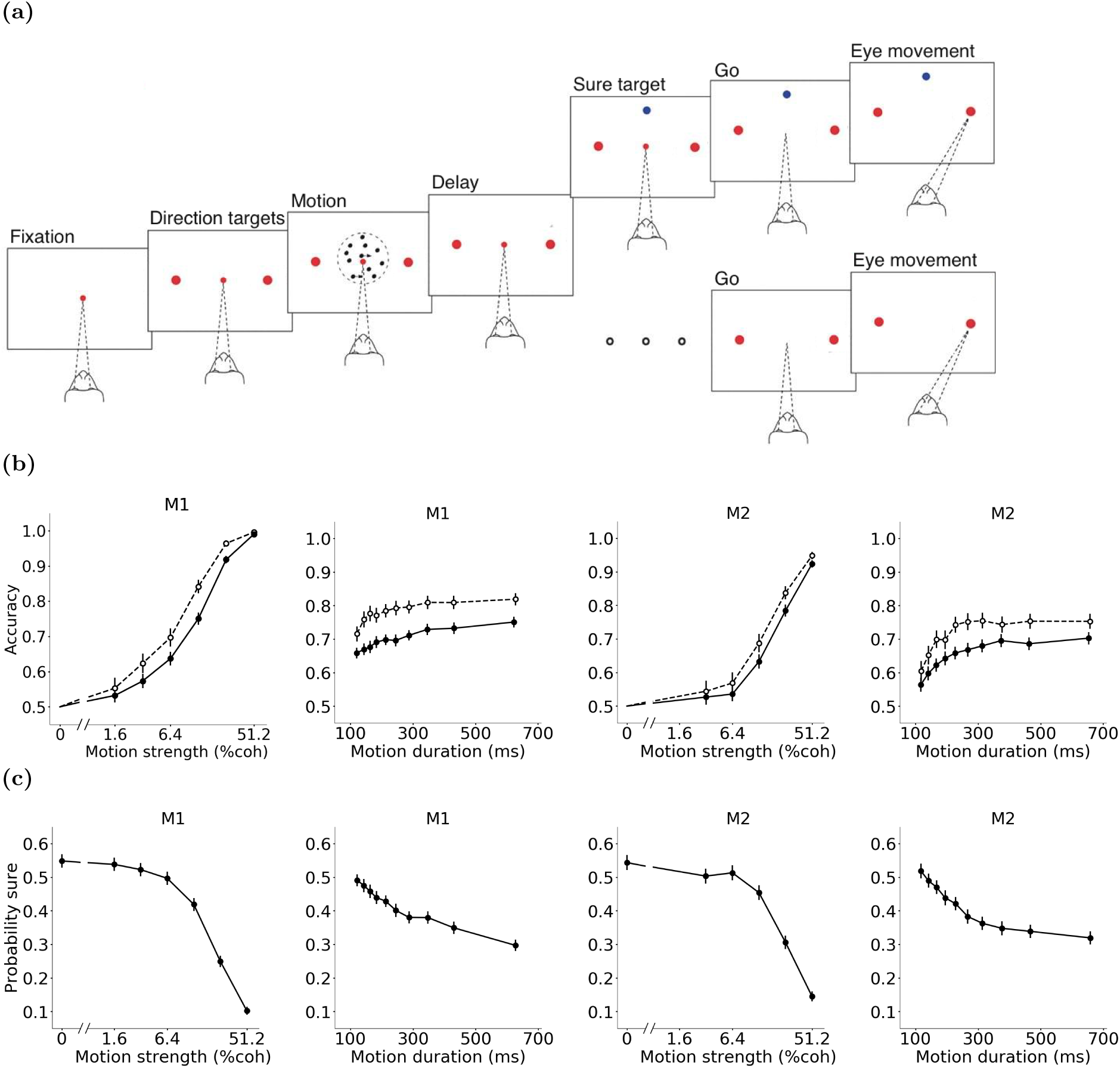
Motion direction discrimination with post-decision wagering. **a)** Task design. On each trial, monkeys viewed a patch of randomly moving dots and made a decision about the net direction of motion. Stimulus strength and duration varied randomly from trial to trial. On half of the trials, only the right and left direction targets were shown (large red dots). The motion stimulus was followed by a delay period. The central fixation point (small red dot) disappeared at the end of the delay (Go cue), instructing the monkey to report perceived motion direction with a saccadic eye movement to one of the two direction targets. Choosing the correct target (right for rightward motion and left for leftward motion) yielded a large reward, whereas choosing the incorrect target resulted in a short timeout. On the other half of the trials, a third target (sure target, shown as a blue dot) appeared on the screen during the delay period. Choosing this target after the Go cue yielded a guaranteed but smaller reward than choosing the correct direction target. **b)** Accuracy as a function of motion strength and duration for the two monkeys (M1 and M2). For the motion strength plots, trials are pooled across all durations. For the motion duration plots, trials are pooled across all strengths. Solid lines show the accuracy on trials where the sure target was not presented. Dashed lines show the accuracy on trials where the sure target was shown but the monkey chose one of the high-stakes direction targets. **c)** Probability of choosing the sure target for different motion strengths and durations for monkeys M1 and M2. Error bars indicate standard error of the mean (s.e.m.).

An optimal decision maker who desires to earn more reward and maximize utility should choose the risky, high-paying direction targets when confident about motion direction and the sure-bet option when doubtful about the correct direction. Monkeys showed a similar behavioral pattern. They chose the sure-bet option more often on more difficult trials, where motion strength was low or motion duration was short (Fig. 1c). Further, when they ignored the sure-bet option and chose the high-stakes direction targets, their accuracy was higher compared to the trials with similar difficulty without the sure-bet option when they had to choose one of the direction targets (trials without sure-bet target; Fig. 1b). These results indicate the presence of a mechanism for assessment of expected decision outcome (confidence), and reliance on this mechanism for guiding the opt-out behavior.

### 2.1 Modeling Perceptual Decision Making with a POMDP

In perceptual decision-making tasks, an ideal observer would infer hidden states of the environment based on a sequence of sensory observations to gain the maximum possible reward utility. This problem can be solved using the general framework of POMDPs, which combines Bayesian inference of hidden states with expected reward maximization (Kaelbling et al., 1998; Pineau et al., 2006; Ross et al., 2008; Rao, 2010; Khalvati and Mackworth, 2013). Formally, a POMDP is a tuple (*S, A, Z, T, O, R*) where *S* and *Z* are two sets containing the states of the environment and observations, respectively. *A* is the set of possible actions. For example, possible actions after each observation in the direction discrimination task are to commit to a choice or to make a new observation. Actions and observations are interleaved with one another: each action results in a new observation. *T* is a transition function that represents the probability of entering a state *s* from a state *s′* after performing an action *a*: *T* (*s, s′, a*) = *P* (*s*|*s′, a*). Note that the environment is assumed to be Markovian, meaning that the next state depends only on the current state and current action. *O* is the observation function, determining the probability of making an observation *z* given a state *s*, i.e., *O*(*s, z*) = *P* (*z*|*s*). The current state of the environment is not known to the decision maker and needs to be inferred based on the history of observations and actions.

A POMDP starts with a prior probability distribution over states of the environment and infers the posterior probability distribution of states after each action and observation. The POMDP model objectively defines belief as this posterior probability. The POMDP belief after *t* observations and actions is given by *b*_*t*_. Starting from an initial belief (*b*_0_), *b*_*t*_ is obtained from *b*_*t−*1_ after action *a*_*t−*1_ and observation *z*_*t*_ as follows:

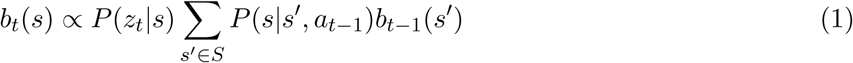

If the actions do not change the state of the environment, *P* (*s*|*s′, a*) = 1 when *s* = *s′* and zero otherwise, and *b*_*t*_(*s*) *∝ P* (*z*_*t*_|*s*)*b*_*t−*1_(*s*). This is usually the case within any trial in most perceptual decision-making experiments, including our direction discrimination task, where the hidden state of the environment is kept fixed.

The POMDP framework also offers objective definitions for the reward function and policy. Each action can potentially yield a reward utility depending on the current state, determined by a reward utility function: *R*(*s*_*t*_, *a*_*t*_). We emphasize reward utility instead of reward size, as the model optimizes the benefit of reward (utility) and the utility of reward does not grow linearly with reward size for a wide range of tasks and behaviors. As the current state is not observable, for each belief and action, there is an expected reward utility, defined as *E*_*s*_*t* [*R*(*s*_*t*_, *a*_*t*_)] = ∑_*s∈S*_ *b*_*t*_(*s*)*R*(*s, a*_*t*_). The optimal decision policy *π*^*∗*^ is a mapping from belief states (probability distributions over states) to actions that maximizes the total expected reward utility (Sondik, 1978):

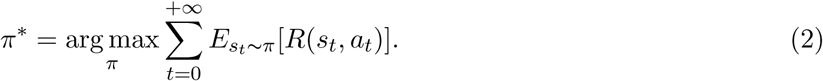

Analogous to previous definitions of confidence as the expected likelihood of success in two-alternative tasks (Kiani and Shadlen, 2009; Kiani et al., 2014; Drugowitsch et al., 2014; Khalvati and Rao, 2015; Hanks and Summerfield, 2017; Pouget et al., 2016; Lak et al., 2017), we define confidence of the POMDP as the expectation of the model that its selected action (choice) is likely to be correct and leads to reward. That is, confidence is the belief of the most probable choice at time *t*.

Similar to how accuracy is calculated for experimental data, choice accuracy for the POMDP model can be defined as the fraction of trials in which the choice corresponding to the highest belief is in fact the correct one and leads to reward:

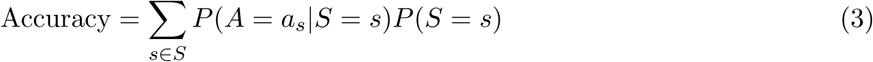

where *a*_*s*_ is the action of choosing state *s*.

Accuracy and confidence could become equal (on average) if the decision maker has an exact model of the environment (Khalvati and Rao, 2015). Such knowledge, however, is unlikely in most natural contexts and common task designs. For example, in the direction discrimination task, subjects face a mixture of stimulus difficulties across trials. They neither know the exact generative function for the stimulus on each trial nor the exact set of motion strengths used in the experiment.

### 2.2 POMDP Model of the Direction Discrimination Task

We first define the hidden state of the direction discrimination task. The random dots stimulus depends on two hidden parameters: direction and coherence. Because motion coherence simply reflects the average strength of motion in a particular direction, we can combine direction and coherence into a single real-valued hidden state for the POMDP model which we call signed motion coherence (Salzman and Newsome, 1994): for the models below, positive values of the signed coherence indicate rightward motion and negative values indicate leftward motion.

The momentary observations *z*_*t*_ at time *t* from a stimulus with signed coherence *c* are modelled as samples drawn from a Gaussian distribution, 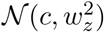, with mean *µ* = *c* and variance 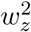 (Fig. 2a). This randomness arises from the stochastic nature of the motion stimulus and sensory neural responses in the motion-selective cortical areas MT and MST (Britten et al., 1992; Celebrini and Newsome, 1994; Kiani et al., 2008).

**Figure 2:**
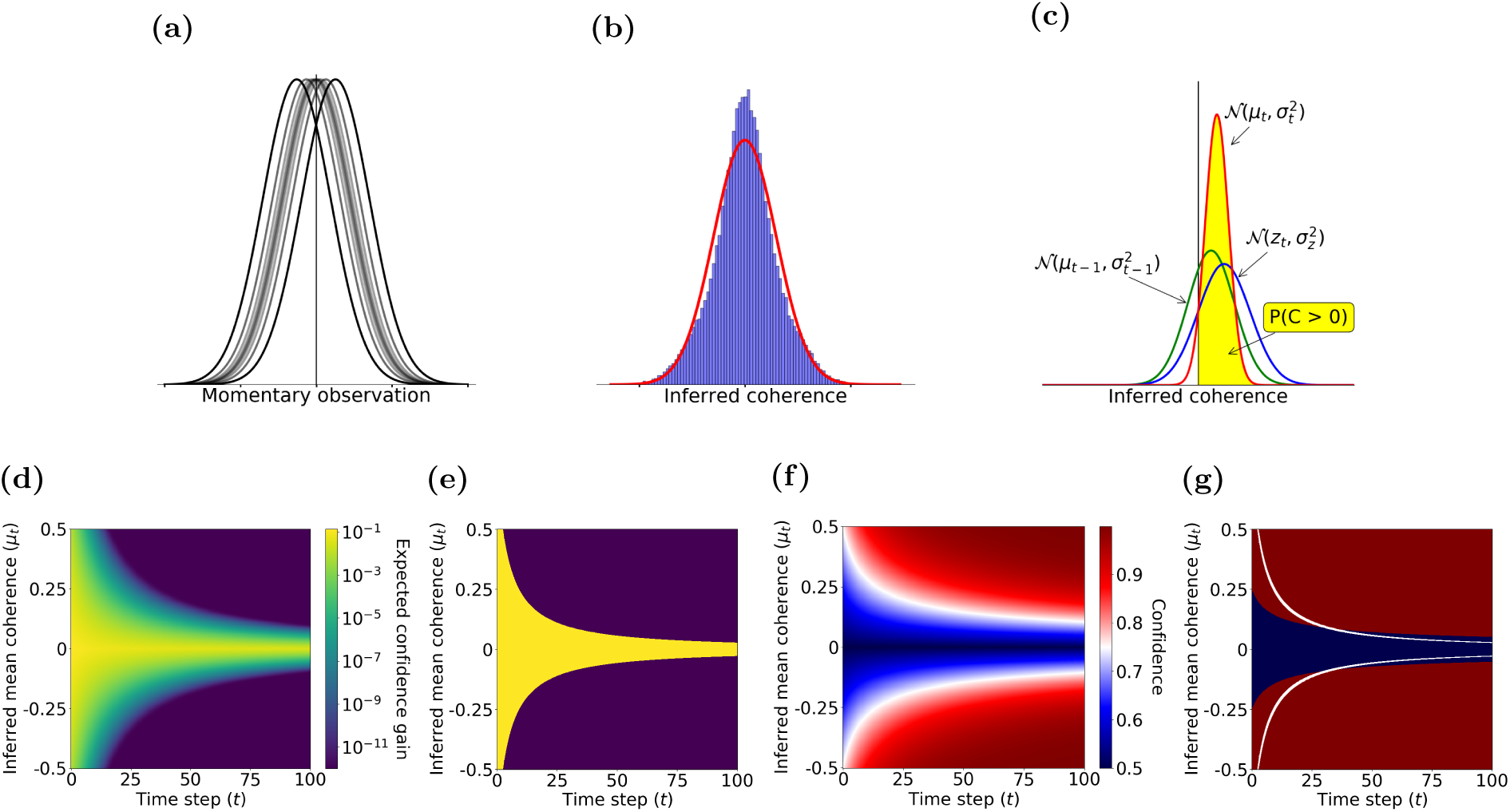
Computation of choice and confidence in the POMDP model of the direction discrimination task. **a)** Probability distribution of momentary observations for a motion coherence *c* is modelled as a Gaussian distribution with mean *µ* = *c* and variance 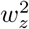. There are multiple Gaussian distributions for different motion coherence. Positive and negative observations indicate rightward and leftward motion directions, respectively. **b)** The distribution of inferred coherence across all trials provides the initial belief state of the POMDP model (blue histogram) at the beginning of each trial. The initial belief is approximated by a Gaussian function (red curve). **c)** The POMDP model sequentially updates its belief about the motion coherence based on new observations. Combining the belief at time *t −* 1 (blue distribution) with the acquired observation from the stimulus at time *t* (green distribution) results in a new belief at *t* (red distribution). The expected likelihood that the rightward choice is correct is the area under the updated belief distribution for positive sensory evidence (yellow region). **d)** The expected confidence gain for a new observation as a function of inferred mean coherence, *µ*_*t*_, and elapsed time, *t*. **e)** An example POMDP decision policy when new observations are associated with a constant cost. The yellow area represents the belief states where the optimal action is to continue observing. The purple area represents the belief states where the POMDP model terminates and commits to a choice. **f)** Confidence as a function of inferred mean coherence, *µ*_*t*_, and time, *t*. **g)** The ratio of reward utilities for sure-bet and correct direction choices determines the POMDP policy for choosing the sure-bet option. The policy can be illustrated as phase boundaries in the confidence plot of panel f. The blue region denotes combinations of inferred coherence and time for which the model would choose the sure-bet target. The red region denotes (*µ*_*t*_, *t*) for which direction targets are chosen. In the simulations in panels d-g, *σ*_*z*_ = 2.0, *σ*_0_ = 1.0, and the utility ratio = 0.63.

The two main actions of our POMDP model are committing to direction *right* or *left*. Also, action “observe” makes the next observation available to update the belief. Finally, the action of choosing the sure-bet option is available during the delay period on half of the trials. The decision maker gets *r*_*right*_ as the reward utility for committing to direction *right* if and only if the direction of the hidden state is *right* (*c >* 0). *r*_*left*_ is the reward utility given to the decision maker by committing to direction *left* if and only if the direction of the hidden state is *left* (*c <* 0). Choosing the sure-bet option, if available, always yields reward utility of *r*_*sure*_.

The POMDP model begins each trial with a prior belief about the signed coherence of the trial. Subjects are not explicitly informed about the exact set of discrete motion coherence levels used in the experiment. They only experience largely overlapping distributions of motion energies on different trials (Kiani et al., 2008). Therefore, it is most realistic to consider that the model’s prior spans a continuous domain, obtained from observations across all trials with various coherence levels and durations. This distribution has a complex shape (Fig. 2b) with a large peak centered at zero. The central peak is due to the high frequency of weak motion coherence in the experiment – a consequence of defining sampled coherence on a log scale (see Methods). Because the logarithmic spacing of motion coherence causes the mass of the prior distribution to be largely concentrated in its central peak, we use a Gaussian approximation to this prior distribution, 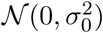. This approximation facilitates developing analytical solutions for our POMDP model. The approximation also matches the notion that humans may be less sensitive to higher order moments than mean and variance (Waskom et al., 2018).

Starting with a Gaussian prior (initial belief), 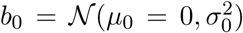, the model iteratively updates its belief about the hidden state of the environment, i.e., the signed motion coherence *c*, following each observation, *z*_*t*_, drawn from the distribution 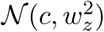 at time step *t* (Fig. 2c). To be able to update the belief, knowledge of the true observation variance, 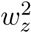, is required. However, 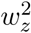 is unknown to the model. Rather, we use 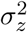 to denote the model’s learned observation variance. This means that the model assumes *z*_*t*_ is drawn from the Gaussian likelihood function 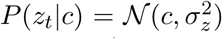. A Gaussian prior and a Gaussian likelihood function together result in a Gaussian posterior (Murphy, 2012) for *c* given by:

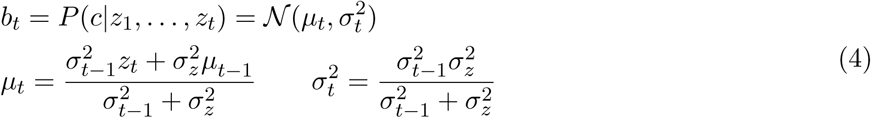

The belief that the motion direction is rightward is the sum of the posterior probability over all positive motion coherence, which is equal to the integral of 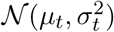 from 0 to +*∞*:

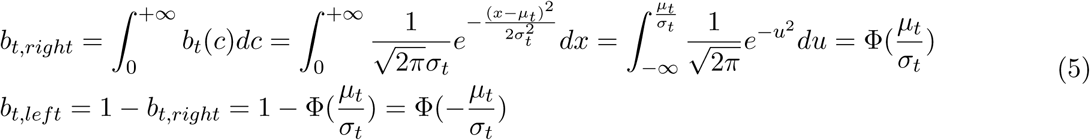

where Φ(*x*) denotes the standard normal cumulative distribution function. For trials where the sure-bet option is not shown, the model chooses the motion direction with the highest belief, that is rightward choice if 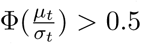 and leftward choice if 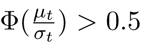. A random choice is made in the unlikely event that 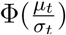 is exactly 0.5. Additionally, as explained in the previous section, the confidence of the model equals the belief about this selected choice:

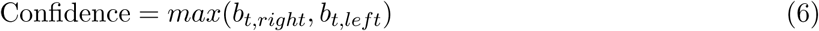

The POMDP model choice accuracy in a trial depends on the likelihood that the motion frames in that trial lead to a stronger belief for the correct motion direction than the incorrect one. This is equivalent to the probability that the inferred mean coherence, *µ*_*t*_, becomes larger than zero for a positive (rightward) motion coherence and less than zero for a negative (leftward) motion coherence. Reorganizing Equation 4 reveals that the sign of *µ*_*t*_ is equal to the sign of the sum of observations 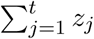 because:

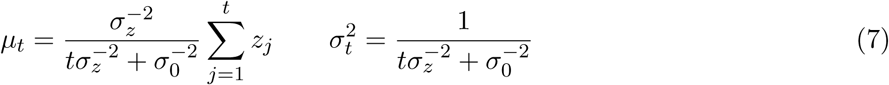

The POMDP approach can easily model termination of the decision-making process and commitment to a choice by assigning a cost (negative utility) to observation gathering and belief update (via the action “observe”)(Rao, 2010). Moreover, because the hidden state does not change with actions within a trial in the motion discrimination task, a one-step look-ahead search (Ross et al., 2008) is adequate to determine the optimal decision policy for non-decreasing observation costs over time (instead of computing the total expected reward utility to the end of the trial; see the proof in Methods). The model halts new observations when the expected increase in confidence is less than the ratio of the cost of an observation and the reward utility for correct choice. The expected increase in confidence after one more observation depends on the current belief and the probability distribution of the next observation according to the model. Specifically, when the current belief is 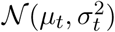, the model assumes that the next observation is a sample from 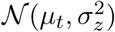, where 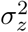 is the learned observation variance. Figure 2d shows the expected increase in confidence for a new observation as a function of two key variables: the inferred *µ*_*t*_ and the elapsed time. The expected increase in confidence from new observations is higher earlier in the trial and for smaller inferred mean coherence, *µ*_*t*_.

A constant observation cost over time, if present, would give rise to a stopping criterion that matches an iso-gain contour. These contours would effectively implement a time-varying bound on *µ*_*t*_ for each motion direction (an upper bound and a lower bound). Figure 2e shows these collapsing bounds for a cost of 10^*−*3^ per observation (in our case, per 10 ms) when the reward utility for a correct direction choice is set to 1. A policy for termination of observations is especially critical in reaction time (RT) tasks where subjects have to decide when to initiate a response. However, a termination policy could exist even in tasks where stimulus duration is controlled by the experimenter, causing early termination of the decision-making process before stimulus offset, especially in long and easy trials (Kiani et al., 2008; Okazawa et al., 2018).

However, if the cost associated with new observations is negligible the model will use all the observations provided in a trial in tasks with experimenter-controlled stimulus durations. In such a case, for a trial with duration *t*, 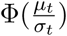 being larger or smaller than 0.5 depends solely on the sign of *µ*_*t*_, and consequently on the sign of the sum of all observations in the trial (see Eq. 7). Therefore, the model accuracy for trials with duration *t* and coherence *c* becomes:

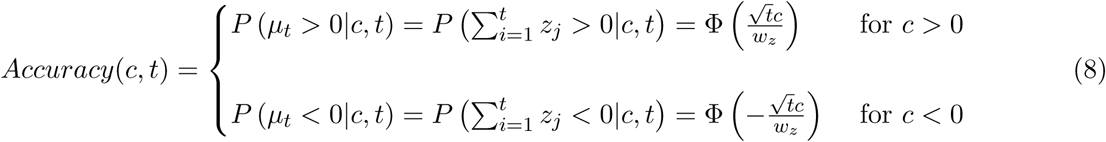

The reward utility maximization principle also determines the choice when the sure-bet option is available. As the reward for the sure-bet option is guaranteed, the POMDP model compares the expected reward utility for choosing each direction with the reward utility for the sure-bet option in order to pick the final action:

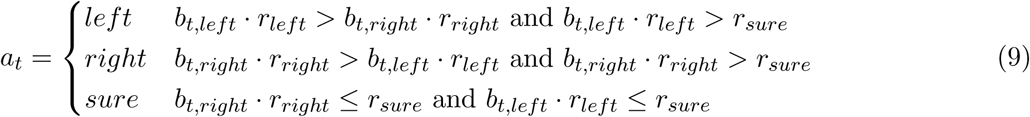

Because *r*_*right*_ = *r*_*left*_ = *r*_*direction*_ in our task, the above policy reduces to a comparison of the confidence with the reward utility ratio of the sure-bet and correct direction choices, *r*_*sure*_*/r*_*direction*_. Since confidence increases with the absolute value of inferred coherence, |*µ*_*t*_|, this reward utility ratio serves as a time-varying boundary that determines the POMDP policy as a function of inferred coherence and time in each direction (upper and lower bounds). Figure 2f shows confidence as a function of inferred coherence and elapsed time for an example POMDP model and Figure 2g shows the model policy for an example reward utility ratio of 0.63.

Equations 4-9 suggest that it is possible to use experimentally measured accuracy in our task to fit the POMDP model parameters. With a constant observation cost, the model has up to four degrees of freedom: (*i*) observation cost; (*ii*) the true observation variance 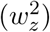, which shapes input samples available to the model; (*iii*) the learned observation variance 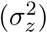, which the model attributes to its inputs; and (*iv*) variance of the prior distribution 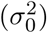. For an optimized POMDP model, however, 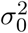 and 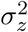 are uniquely determined by 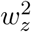 and observation cost. As mentioned before, 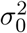 determines the prior belief, which should be consistent with the overall distribution of states and consequently, perceived observations. Moreover, 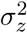 should match the model’s posterior belief with its average accuracy for each motion duration. This is possible based on the feedback given about motion direction choices (correct or wrong) after each trial (see Methods for details of obtaining these parameters). Such a model, therefore, has two degrees of freedom: observation cost and 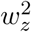.

Note that correct posterior belief (matched with accuracy on average) is not necessary for maximizing the reward utility in choosing between the two directions because determining the sign of the sum of observations is sufficient. However, it is necessary for the wagering task where the expected reward utility of choices needs to be computed (Eq. 9).

### 2.3 Comparison of Model Predictions with Experimental Data

In our task, the stimulus viewing duration was controlled by the experimenter and subjects were required to maintain fixation throughout the duration. As a result, the cost of acquiring new observations while maintaining fixation on the stimulus could be negligible. We verified this hypothesis by comparing the model with two degrees of freedom (observation cost and *w*_*z*_) to a POMDP that uses all observations in each trial (Equation 8 with only *w*_*z*_ as the free parameter). They were not significantly different in quality of fits even without penalizing the extra free parameter (Vuong’s closeness test (Vuong, 1989), *p* = 0.16 for monkey M1 and *p* = 0.07 for monkey M2; see Methods).

Therefore, using Equation 8, we fit the model to each monkey’s accuracy on trials in which the sure target was not shown (Fig. 3a) (*R*^2^ = .95 and .88 for monkeys 1 and 2, respectively) and obtained the observation variance 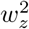. We then used the estimated 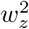 to construct the variance of the prior belief, 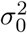 (Fig. 2b). Finally, we used the estimated 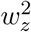 and prior belief to estimate the learned observation noise, 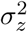, by fitting the model belief about the choice to the accuracy on trials without the sure-bet option (See Methods for details of this fitting process).

**Figure 3:**
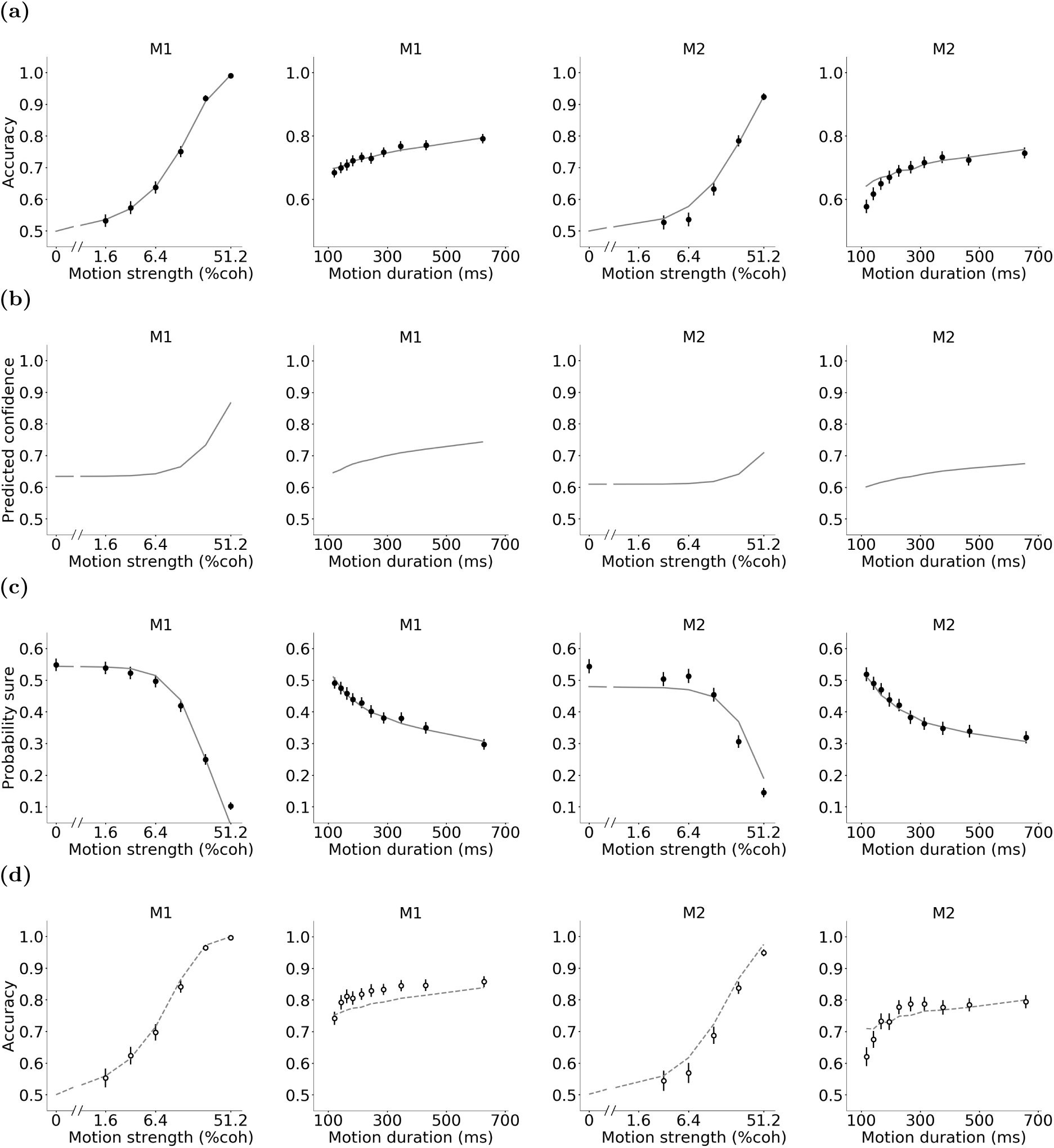
The POMDP model captures the monkey’s behavior. **a)** The model was fit to each monkey’s accuracy on trials without the sure-bet option. Solid lines are model fits and data points are the measured accuracy for each motion strength and duration for monkeys M1 and M2. **b)** The model parameters obtained from the fits in (a) were used to predict confidence for each motion strength and duration for each monkey. **c)** Predictions of the POMDP model about confidence were thresholded to fit the likelihood of choosing the sure-bet option (see Methods). **d)** With the model parameters fully constrained by accuracy on trials without the sure-bet target and the likelihood of choosing the sure-bet target when it was presented to the monkey, we predicted the monkey’s accuracy on trials in which the sure-bet target was shown but ignored. Lines are model predictions. Data points are identical to those in Figure 1. Error bars indicate s.e.m.

Having estimated the model parameters based on trials without the sure-bet target, we predicted the monkey’s confidence for each motion coherence and duration (Fig. 3b). These predictions suggested a monotonic increase in confidence with motion coherence and duration, compatible with previous studies (Vickers, 1979; Balakrishnan and Ratcliff, 1996; Kiani and Shadlen, 2009; Hangya et al., 2016; Zylberberg et al., 2016).

Since the model chooses the sure-bet option when confidence (belief) is less than the reward utility ratio of the sure-bet and correct direction choice (Eq. 9), it predicts lower probability of choosing the sure-bet target on trials with stronger motion and longer durations. Since we do not know the exact utility of reward volumes associated with the sure-bet and correct direction choices, we added a new free parameter to the model that represented the reward utility ratio and used this parameter as a threshold that the confidence was compared to on trials in which the sure-bet target was presented. Optimizing this parameter in order to match the predicted confidence of the POMDP model with the monkey’s behavior provided a fit with *R*^2^ = 0.90 and 0.82 for monkey M1 and monkey M2, respectively (Fig. 3c).

Since the model parameters are fully specified based on the monkey’s accuracy on trials without the sure-bet target and the probability of choosing the sure-bet target when it was presented, we could provide quantitative predictions for the monkey’s direction choice accuracy when the sure-bet target was presented but not chosen. Figure 3d shows these predictions (gray dashed lines), demonstrating that they closely matched experimentally measured accuracy on trials where the monkey ignored the sure-bet option (*R*^2^ = 0.90 and 0.81 for monkey M1 and monkey M2, respectively).

### 2.4 POMDP Model Predicts Experimentally Observed Discrepancies between Accuracy and Confidence

Because our POMDP model enables us to predict confidence from accuracy, we explored if it could also explain five well-documented discrepancies between accuracy and confidence. Based on the model’s success, we suggest that these discrepancies are neither anomalies of the decision-making process nor do they necessarily indicate a divergence of the neural mechanisms that compute choice and confidence. Rather, these phenomena are expected signatures of a decision-making process that infers the choice and its associated confidence in a unified framework.

#### 2.4.1 Hard-easy effect

The hard-easy effect, which has been documented extensively (Juslin et al., 1997; Sanders et al., 2016), is the tendency to overestimate the likelihood of one’s success for difficult decisions and underestimate it for easy decisions. This effect emerges naturally in our POMDP model, as illustrated in Figure 4a. In the face of uncertainty about the hidden state, the model computes confidence over all possible stimulus strengths. The uncertainty about the stimulus strength, therefore, causes overconfidence in difficult trials and underconfidence in easy ones.

**Figure 4:**
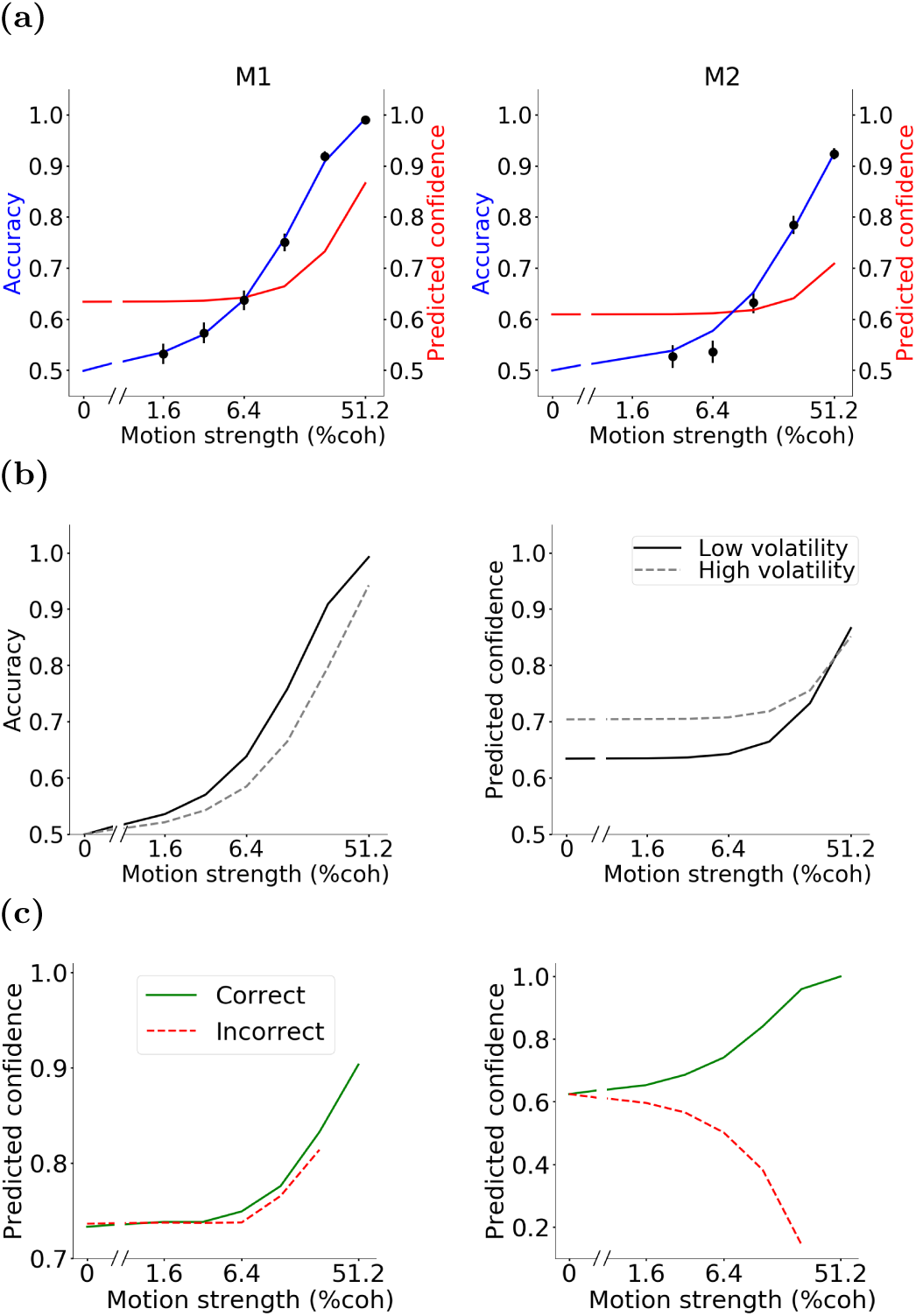
POMDP model explains systematic deviations of confidence reports from accuracy, experimental manipulations that dissociate accuracy and confidence, and sensitivity of confidence reports to experimental methods for measuring confidence. Here, we highlight three commonly-observed experimental effects. **a)** The hard-easy effect. Uncertainty about the strength of observed evidence makes the model more confident than warranted by accuracy on hard trials and less confident than warranted by accuracy on easy trials (compare blue and red curves). The psychometric and predicted confidence functions are adopted from Figures 3a and 3b. **b)** The volatility of observations has opposing effects on accuracy and confidence. Artificial increase of volatility following training with lower volatility reduces accuracy (left panel) but boosts confidence (right panel). This dissociation occurs because the model relies on the observation noise learned during the lower-volatility training to render choices on the higher volatility trials. **c)** The model is sensitive to simultaneous vs. sequential reports of choice and confidence in reaction-time tasks. Simultaneous reports are commonly associated with higher confidence for erroneous choices on trials with stronger stimuli (left panel). This pattern is partly caused by lower decision times for stronger stimuli and the dependence of model confidence on elapsed time (Fig. 2f). In contrast, sequential reports of choice and confidence could be associated with reduced confidence for incorrect choices for stronger stimuli (right panel). This pattern is caused because sensory and motor delays render the last observations inconsequential for the choice but the model uses those observations to refine its confidence report following the choice. The discrepancy between the left and right panels therefore arises because confidence reports in the sequential task designs are informed by more observations than used for the choice.

#### 2.4.2 Opposing effects of the volatility of observations on choice and confidence

A common observation in past studies has been that increasing the variability of the stimulus reduces subjects’ accuracy but increases their confidence about their choices (Rahnev et al., 2011; Fetsch et al., 2014; Zylberberg et al., 2016). Our POMDP model shows that this seemingly paradoxical effect of stimulus variability arises naturally in an optimal inference framework when the subject does not have access to the true model of the environment (in this case, the true observation noise).

The stimulus variability effects have been explored in tasks where subjects were trained using a baseline (lower) stimulus variability, before being tested with higher variability. Further, trials with different levels of stimulus variability were randomly intermixed. Consequently, our model postulates that subjects continued to rely on the observation noise learned during initial training, and used this model for choice and confidence on the high volatility trials. Higher volatility (larger 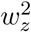) decreases accuracy (Fig. 4b, left plot) by generating more overlapping observations from different motion directions (Eq. 8). Higher volatility also generates extreme observations (far from the mean) more often. This higher frequency of extreme observations is not expected based on the observation noise learned during training 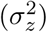. As a result, the POMDP framework considers the extreme observations highly discriminative, resulting in higher confidence (Fig. 4b, right plot), as observed in past experiments.

#### 2.4.3 Sensitivity of confidence measurements to simultaneous versus sequential reports of choice and confidence

The POMDP model can also be applied to reaction time tasks (besides fixed duration tasks), where subjects report their choice as soon as they are ready. In these tasks, the experimenter may ask for either a simultaneous or sequential report of choice and confidence (Kiani et al., 2014; Sanders et al., 2016; van Den Berg et al., 2016). A simultaneous report of choice and confidence ensures that both are based on the same observations about the stimulus (Kiani et al., 2014). In contrast, a sequential report of choice and confidence opens up the possibility of revising confidence based on the last few sensory observations not used to make the choice (van Den Berg et al., 2016). The sensory and motor delays in the neural circuits underlying decisions usually amount to around 250 ms or more, implying that the last 250 ms of the stimulus do not contribute to the choice. Observations made during this time period, however, can contribute to a change of mind (Resulaj et al., 2009; Kiani et al., 2014) or change in confidence (van Den Berg et al., 2016) after the choice is made. Confidence on error trials is especially susceptible to such revisions.

Past experiments have shown that for simultaneous reports of choice and confidence, confidence for incorrect choices often increases with stimulus strength, compatible with the predictions of bounded accumulation models such as the drift diffusion model (DDM) (Kiani et al., 2014; van Den Berg et al., 2016). However, for sequential reports, confidence can be revised based on the last parts of the stimulus not used for the reported choice, causing confidence for incorrect choices to decrease with stimulus strength, compatible with the predictions of signal detection theory (Kepecs et al., 2008; Sanders et al., 2016; van Den Berg et al., 2016).

A reaction time version of the POMDP model reproduces both of these results as shown in Figure 4c. For simultaneous reports, confidence increases with stimulus strengths for both correct and incorrect responses as higher stimulus strengths tend to have shorter reaction times and therefore higher confidence compared to lower stimulus strengths (left panel). However, introducing a delay between choice and the confidence report and allowing confidence to be revised based on additional observations after the choice is made reverses the confidence trend in incorrect trials (right panel). Note that when confidence is reported after a choice, it can become less than 0.5 due to a change of mind about the correct choice.

#### 2.4.4 Discrepancy of sensitivity for accuracy (*d′*) and confidence (meta-*d′*)

The POMDP model also explains experimentally observed differences between the sensitivity of accuracy and confidence to observations, commonly quantified with *d′* and meta-*d′*, respectively (Maniscalco and Lau, 2012). *d′* and meta-*d′* are defined based on a signal detection theory (SDT) framework. *d′* quantifies the difference of sensory evidence distributions underlying the probability of correct and incorrect choices. However, meta-*d′* is related to the distribution of confidence ratings for those choices. For a binary confidence rating (low or high confidence, similar to rejecting or choosing the sure-bet option), meta-*d′* contrasts the probability of a high confidence rating for a correct response with that of an error. Some studies have reported that confidence ratings are not consistent with the sensitivity of the choice accuracy (*d′*) (Maniscalco and Lau, 2012; Fleming and Dolan, 2012; Charles et al., 2013; Palmer et al., 2014). However, for an SDT ideal observer meta-*d′* and *d′* have to be similar in the absence of variability in the confidence rating threshold (Fig. 5a). Therefore, it has been suggested that the different meta-*d′* and *d′* in experimental data must be due to loss of information for confidence judgments or different neural mechanisms for confidence and choice (Maniscalco and Lau, 2012; Fleming and Daw, 2017).

**Figure 5:**
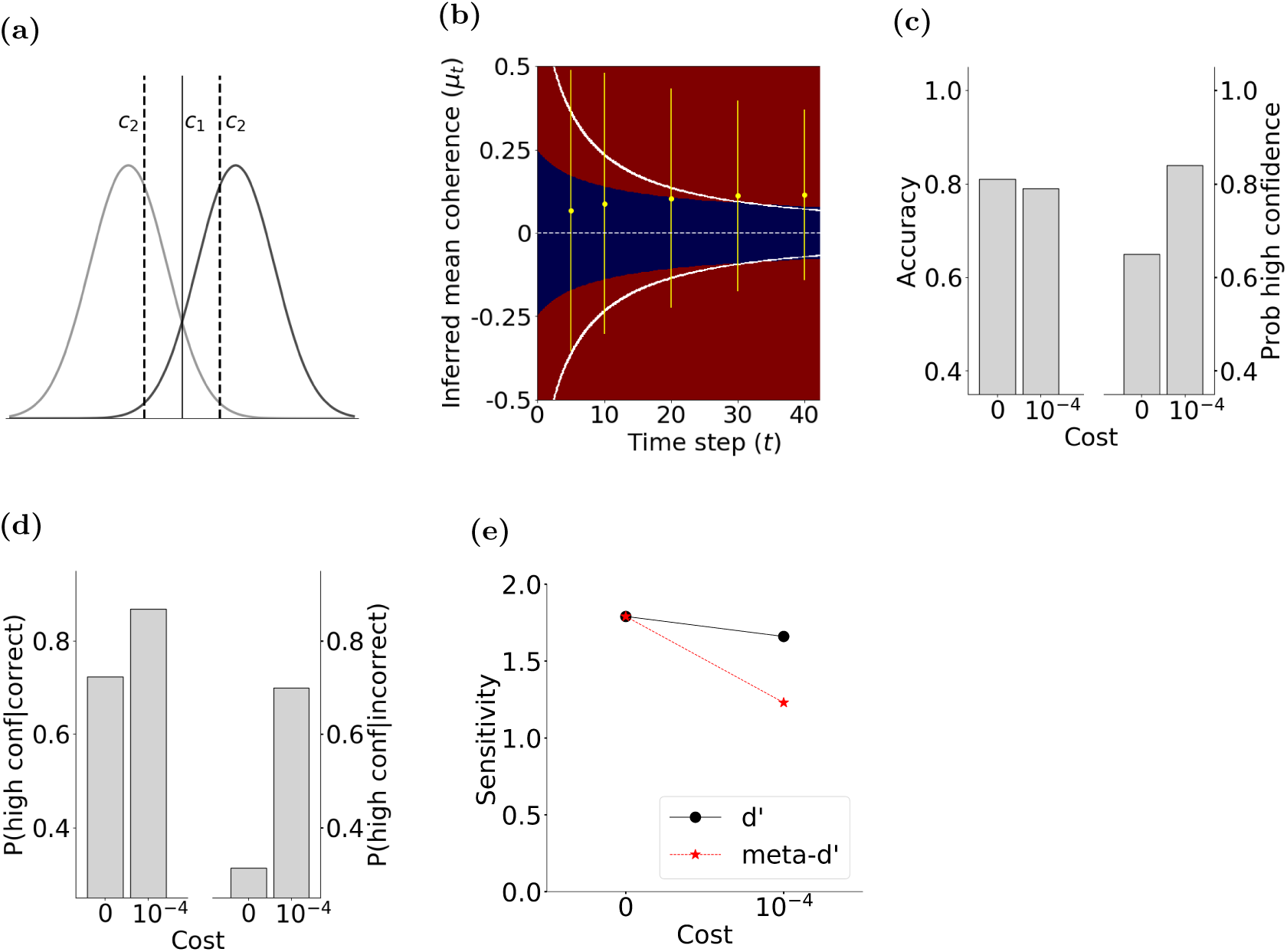
POMDP model explains different values for *d′* and meta-*d′*. **a)** A signal detection theory framework predicts identical *d′* and meta-*d′* when the same observations inform both choice and confidence rating. The competing stimuli (e.g., right and leftward motion of a particular coherence) give rise to two observation distributions. The *c*_1_ criterion is used for choosing right and left, and the *c*_2_ criteria are used to report low or high confidence for each choice. *d′* quantifies type 1 sensitivity: the distance between the distributions in units of standard deviation. Meta-*d′* quantifies type 2 sensitivity: the separation of the two distributions compatible with hit rate and false alarm rate of confidence reports: *p* (high conf|correct) and *p* (high conf|incorrect), respectively. In SDT, *d′* and *c*_1_ fully constrain meta-*d′* and an optimal meta-cognitive observer must have equal *d′* and meta-*d′* (Maniscalco and Lau, 2012). **b)** Observation noise could cause highly variable *µ*_*t*_ at the beginning of a trial, and thus temporarily produce excessive confidence. This excessive confidence may become permanent if the decision-making process is stopped by reaching the termination bounds. Solid white lines show the two decision termination bounds (observation cost, 10^*−4*^). Thresholds for separating low and high confidence ratings are shown as boundaries between blue (low confidence) and red (high confidence) regions. The horizontal dashed line shows the boundary that separates right and left direction choices based on the sign of the inferred coherence. Yellow dots and lines show *mean±*2 *× s*.*d*. (95% of the distribution mass) of the inferred coherence for a particular stimulus strength (c = +12.8%) at a few different time steps (10 ms per step). Temporary excessive confidence due to early termination is more prominent for the incorrect trials (negative *µ*_*t*_ in this simulation). **c)** Early termination can cause a modest reduction of accuracy and a marked increase of high-confidence ratings. **d)** Early termination can cause a larger increase in the probability of high confidence ratings for incorrect than correct choices. **e)** Changes of accuracy (c) and confidence ratings (d) can lead to a larger drop in meta-*d′* than *d′*. Model parameters are identical to those for monkey M1, except for the observation cost.

In the absence of an observation cost, where the POMDP model uses all available evidence, its *d′* and meta-*d′* match each other, similar to SDT. That would be true regardless of whether the decision maker does or does not have access to the exact model of the environment. However, if there is an early termination of information gathering, then meta-*d′* could diverge from *d′*. This discrepancy emerges in the model not because of distinct mechanisms for choice and confidence, but because early terminations of the decision-making process have quantitatively distinct effects on the choice accuracy and the likelihood of high confidence ratings for correct and incorrect choices. Because early terminations curtail the use of evidence, they reduce accuracy and, therefore, decrease *d′*. Further, in the face of uncertainty about the reliability of evidence, early terminations are associated with higher confidence (Figs. 2f and 5b). This combination means that for a wide range of model parameter values, the model makes more errors but it is also more confident about its choices compared to a model without an observation cost. Critically, the confidence is inflated more on error than correct trials (Fig. 5d, reducing meta-*d′*. This reduction could be substantially larger than the reduction of *d′*. Consequently, the model could generate meta-*d′* values smaller than its *d′*, even though it computes the choice and confidence through the same optimal process.

Figure 5 illustrates these effects by simulating intermediate coherence (+12.8%) trials with 400 ms duration and subjecting the model choices and confidence to the *d′* and meta-*d′* calculations. Model parameters are inherited from Monkey M1 except for the addition of an observation cost (10^*−*4^/observation). Early in the trial, observation noise can temporarily produce large positive or negative inferred *µ*_*t*_, and thus high confidence (Fig. 5b, yellow lines illustrate *mean ±* 2 *× s*.*d*. of the inferred *µ*_*t*_). Such large *µ*_*t*_ are much less likely at later times because of the correction of excessive early confidence with additional observations. These later corrections, however, are prevented if the termination bounds (Fig. 5b, white lines) are reached earlier. Such occasional early terminations reduce the model accuracy by only 2% for this motion coherence (from 81% with no observation cost to 79%), but increase the overall probability of high confidence choices by 19% (from 65% to 84%) (Fig. 5c). The corrective effect of additional observations on confidence is more pronounced when the initial choice is incorrect as new observations are more likely to cancel the extreme noise that lead to early error choices. Consequently, early terminations increase the fraction of high confidence responses for incorrect choices by 39% (from 31% to 70%), whereas the increase for correct choices is 15% (from 72% to 87%) (Fig. 5d). This reduces the contrast of confidence for correct and error choices, resulting in a reduction of meta-*d′*. This reduction is larger than the very modest reduction of *d′*, bringing the ratio of meta-*d′* to *d′* to 0.74, significantly below 1 (Fig. 5e).

The reduction of meta-*d′* could happen even when the overall confidence rating does not increase in the model, as meta-*d′* depends on the contrast of confidence for correct and error choices, which could be differentially affected by early terminations with or without an overall confidence increase. Generally, meta-*d′* to *d′* ratios below 1 are common for a wide range of POMDP model parameters matching a common result in past behavioral studies (Fleming and Daw, 2017). Further, the model predicts a mismatch between meta-*d′* and *d′* in reaction-time tasks, where the decision maker initiates a response as soon as reaching a decision. Overall, distinct *d′* and meta-*d′* values can arise in the POMDP framework not because different information shapes choice and confidence, but rather when the decision-making process can stop due to a termination criterion without utilizing all the available information. This important alternative has been largely neglected in past explanations of mismatching meta-*d′* and *d′*.

#### 2.4.5 Effects of choice-congruent and choice-incongruent evidence

The last phenomenon we explore in this section is whether confidence reports are more strongly influenced by evidence congruent with the choice compared to incongruent evidence. Previous studies have reported that whereas choice is shaped by the balance of evidence for different options, confidence is more strongly shaped by choice-congruent evidence (Zylberberg et al., 2012; Peters et al., 2017). These results have been interpreted as support for processes that compute confidence after the choice by readjusting the weight of evidence based on the choice (a form of confirmation bias). Our POMDP model demonstrates that this interpretation is not unique. Rather, existing experimental results could be explained without assuming distinct choice and confidence processes, or choice-dependent re-weighting of evidence.

A key feature of analyzing data based on the POMDP framework is to distinguish the observations used by the subject and those analyzed by an experimenter who monitors the subject’s behavior. Because the experimenter does not have access to the subject’s observations as encoded in the nervous system, analysis of data has to rely on the expected distribution of evidence given stimulus properties (e.g., using filters on the stimulus (Zylberberg et al., 2012)) or recordings from the brain (e.g., electrocorticography or ECoG (Peters et al., 2017)). Such estimated observations could markedly diverge from the actual observations used by the subject. A wide variety of mechanisms could underlie such a divergence, including decision bounds or other termination criteria unknown to the experimenter, sampling rates that mismatch the stimulus design, shifts in spatial or temporal attention during a trial, noise in the representation of sensory information by neural responses, or recording noise from the brain.

To clarify the significance of the divergence of observations used by the decision maker and those the experimenter uses to investigate behavior, consider the case where a decision maker uses only a proportion of the observations analyzed by the experimenter (*n* out of the total *t* samples, *n < t*). In this situation, the *t − n* samples not used by the subject act as noise in the analyses. Classification of choice based on stimulus fluctuations reveals equal and opposing influence of stimuli supporting different alternatives as both used and unused observations come from the same distribution. However, conditional on the subject’s choice, the proportion of choice-congruent observations is higher in the portion of the stimulus used by the subject, compared to the unused portion. This is simply because the sum of random variables drawn independently from the same distribution being positive is evidence in favor of each of these variables being positive. If we reorder the observations in a way that *z*_1_, …, *z*_*n*_ become the ones used by the subject and *z*_*n*+1_, …, *z*_*t*_ are the unused ones (only to simplify the equations), we have:

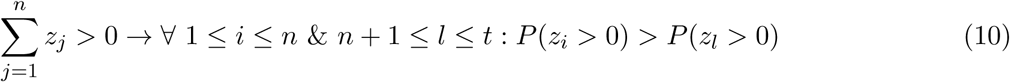

And also:

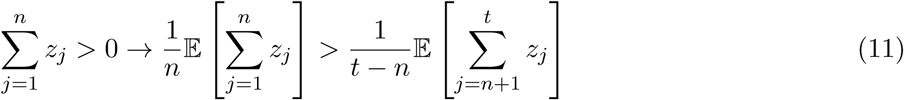

The inequalities of Equations 10 and 11 have a profound side effect for quantification of the influence of individual observations on confidence. If we divide observations based on whether they support the choice, the ratio of total choice-congruent observations to total incongruent observations will be higher for the set of observations used by the decision maker (the *n* samples) than those used in the experimenter’s analyses (all *t* samples). As a result, a classifier that uses all *t* observations to predict the decision maker’s confidence has to give a larger weight to the choice-congruent observations to compensate for the dilution of congruent evidence caused by the unused stimulus samples.

To demonstrate this, we simulated a fixed-duration version of the random dots task with binary confidence ratings (low vs. high). For any stimulus strength and with *n < t*, a logistic classifier fit to the proportion of high confidence ratings by the POMDP model yielded larger weights for congruent than incongruent observations (Fig. 6a). In contrast, a logistic classifier fit to right and left choices based on stimulus fluctuations revealed equal and opposing weights for positive and negative samples as both used and unused observations come from the same distribution. As expected from Equation 11, the imbalance of the weights of the confidence was more pronounced for smaller *n*. To further demonstrate the inevitable imbalance of the weights, we compared the prediction accuracy of the confidence classifier with two alternative classifiers: one forced to have balanced weights for congruent and incongruent observations and a second classifier that had access only to the congruent evidence (Fig. 6b). Similar comparisons were used in past studies (Peters et al., 2017). The confidence classifier with balanced weights had a lower prediction accuracy, especially for low *n*, where its accuracy was even lower than the classifier that totally ignored incongruent observations.

**Figure 6:**
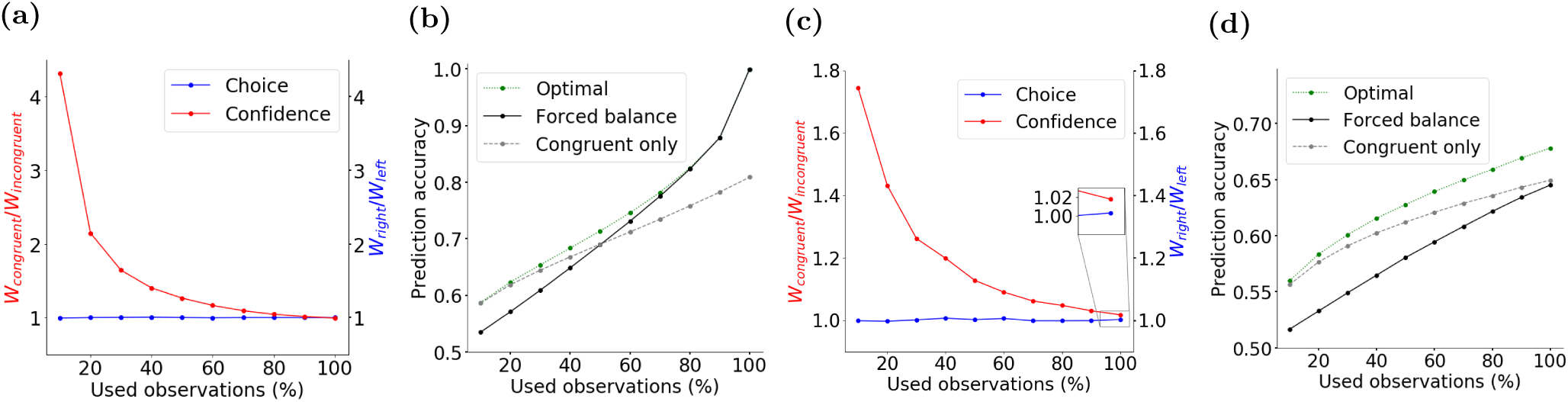
POMDP model explains seemingly higher influence of choice-congruent evidence on confidence ratings. Discrepancy in the observations used by a decision maker and those used by an experimenter studying the decision maker’s behavior could lead to biased interpretation of experimental results. **a)** We simulated a POMDP model that uses a fraction of observations available in a trial unbeknownst to the experimenter. The observations supporting opposing choices equally inform the model’s behavior (Eqs. 4-9). However, an experimenter who uses a classifier to predict choices and confidence based on all observations in the trial finds an apparently larger influence of choice-congruent observations on confidence. **b)** Forcing the classifier to have balanced weights for all observations causes lower prediction accuracy of confidence ratings, especially when the proportion of used evidence is low. In such cases, even a model that totally ignores choice-incongruent observations performs better than the balanced model. However, the better performance of models with imbalanced weights does not reflect the decision making process. It stems merely from the experimenter’s lack of knowledge about the observations used by the simulated model.**c-d)** Same as a-b but observations accessible to the experimenter are noisy estimates of observations available to the decision maker. Such noise reduces the prediction accuracy of the experimenter’s classifier, but more important, it also causes imbalanced weights in the optimal classifier (c) and lower performance of the balanced classifier (d). That is true even when both the decision maker and experimenter use all the available observations (inset in c). The noise in these simulations comes from a zero-mean Gaussian distribution with a variance of 25% larger than 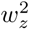.

With similar reasoning, choice-congruent observations gain a higher weight in predicting confidence when the experimenter uses a subset of the observations used by the subject (Supplementary Fig. 1).

Although the example above focuses on a particular source of discrepancy between the observations used by the decision maker and experimenter (different number of used samples), the conclusion generalizes to other sources of discrepancy. Some of these sources such as neural noise are almost always present and quite difficult to correct the analyses for. Essentially, the observations analyzed by the experimenter are almost always noisy estimates of the observation used by the decision maker: 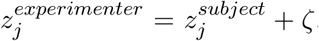, where *ζ* denotes noise with an often unknown magnitude. The neural noise causes the same dilution of choice congruent evidence explained in the example above. Consequently, the experimenter is bound to estimate a higher weight for congruent samples in the analyses even when *n* = *t* and even though such weight imbalance may not exist for the decision maker. Large enough noise can even make a classifier that only uses choice-congruent observations better than a balanced classifier (Figs. 6c and 6d).

### 2.5 Relationship between POMDP and Drift Diffusion Model

A simple mathematical model that has been extensively used to provide quantitative fits to behavior and explain neural activity in various brain regions is the drift diffusion model (DDM) (Ratcliff et al., 2016). DDM assumes that each observation confers evidence in favor of one choice and an equal amount of evidence against the other choice (Fig. 7a). Integration of sensory evidence over time provides a decision variable (DV) that tracks the total evidence in favor of each choice. In most formulations of DDM, two bounds above and below the initial value of the DV (+B and -B in Fig. 7a) act as termination criteria for the decision. As soon as the DV reaches one of these bounds, the decision-making process stops and the choice associated with the bound is made. In cases where the stimulus terminates before a bound is reached, the choice with the most supporting evidence is selected (Kiani et al., 2008). For the direction discrimination task, the decision variable, *V*_*t*_, is updated with each new sensory observation according to:

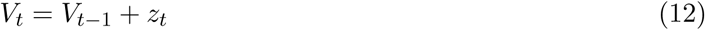

where *V*_*t−*1_ is the DV at time *t −* 1 and *z*_*t*_ represents the momentary sensory observation drawn from a Gaussian distribution with mean *c* and variance 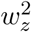. *V*_0_ is initialized to zero when the prior probability and expected reward of the two choices are equal. Therefore, prior to reaching a bound, *V*_*t*_ equals the sum of observations 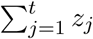 at time *t*.

**Figure 7:**
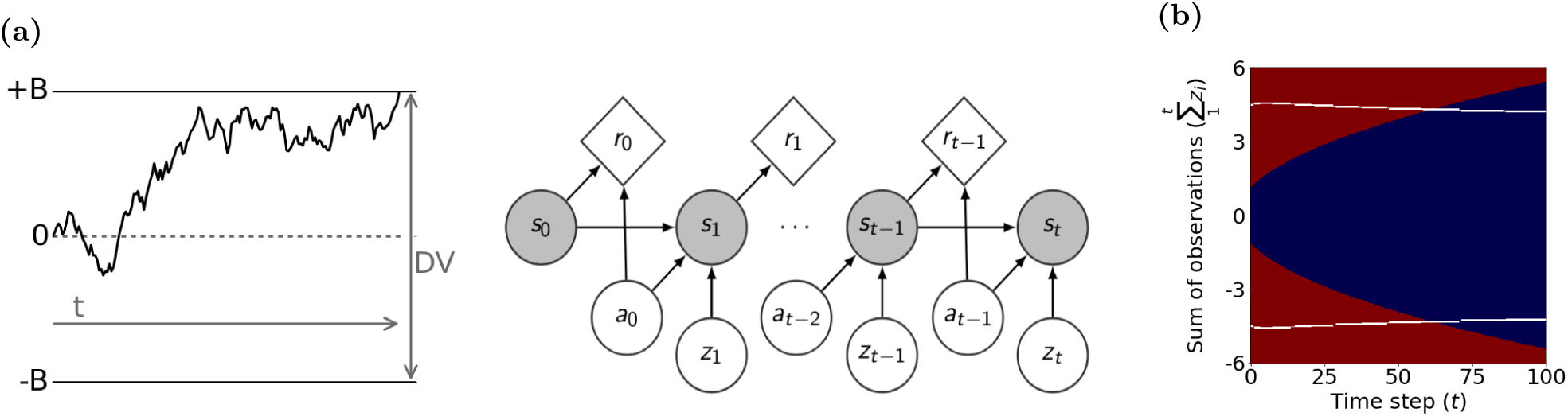
The POMDP policy can be implemented by a drift diffusion model (DDM) with collapsing bounds. **a)** In the standard DDM, the decision variable (DV) is the sum of observations over time. The process stops when the DV reaches one of the static decision bounds (+B or -B). The POMDP model, however, infers a posterior probability distribution over the hidden state of the environment based on observations. It commits to a choice when the expected increase in the probability of a correct response is not worth the cost of an extra observation. **b)** The time-varying bounds on *µ*_*t*_ in the POMDP policy map (e.g., solid white line in Fig. 5b) have equivalent time-varying bounds on the DV in the DDM (Eq. 14; white lines in this panel). Similarly, the low and high confidence regions of the POMDP policy map have equivalents in the DDM.

Although the POMDP model uses Bayes rule for updating the mean, *µ*_*t*_, and variance, 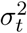, of the belief function, its update rule can be mapped to to the DDM. Taking the ratio of the two update rules in Equation 7, we obtain:

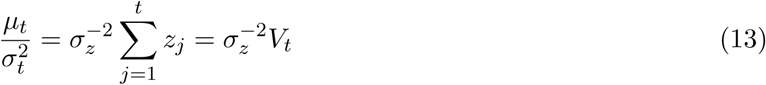

where the second equality is based on the definition of the DV in DDM (Eq. 12). Thus, the Bayes update of the inferred coherence, *µ*_*t*_, can be achieved via addition in the DDM. This means that there is a unique mapping from *µ*_*t*_ and 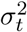 of a POMDP model to the *V*_*t*_ and *t* of the DDM.

This mapping holds in the presence of a bound in the DDM (Moreno-Bote, 2010; Drugowitsch et al., 2014). Moreover, the termination criterion in the POMDP model translates to a unique bound in the DDM. As shown in Figures 2e and 5b, the policy in the POMDP model can be expressed as a bound in the space defined by inferred mean coherence, *µ*_*t*_, over time. This bound on *µ*_*t*_ has an equivalent bound on *V*_*t*_ in the DDM. In general, if Θ*′* (*t*) is the time-varying termination criterion applied to *µ*_*t*_ in a POMDP model (as in Fig. 2e), the equivalent bound, Θ (*t*), on *V*_*t*_ in the DDM is given by:

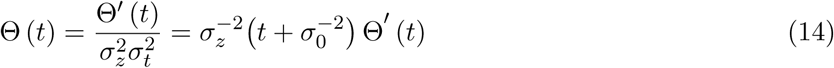

where the first equality derives from Equation 13 and the second from Equation 7. Similarly, confidence ratings can be expressed as time-varying boundaries in the DDM. Figure 7b shows the decision bound and confidence rating boundaries based on the accumulated evidence in the DDM derived to match the POMDP model in Figure 5b.

Overall, both the inference process and the termination criterion of the POMDP model can be implemented with a DDM, suggesting that the neural circuitry for integration of sensory evidence could effectively be implementing the POMDP policy explained in this paper.

## 3 Discussion

We present a Bayesian framework based on partially observable Markov decision processes (POMDPs) that accounts for choice and confidence in perceptual decision-making tasks. Our framework explains the effects of observation cost and task structure on choice and confidence. It also elucidates how the observation noise learned by a Bayesian decision maker may systematically differ from the veridical observation noise, and how this difference influences prior beliefs and confidence. We use our framework to explain the emergence of commonly-observed discrepancies between confidence and choice accuracy. Further, we show how our POMDP model can be mapped to bounded evidence accumulation models (Link, 1992; Vickers, 1979; Ratcliff et al., 2016) and potentially implemented by the same cortical and sub-cortical neural networks implicated in the decision-making process (Gold and Shadlen, 2007; Schall, 2003).

We tested our model using the behavioral data of monkeys performing a direction discrimination task with post-decision wagering (Kiani and Shadlen, 2009). The monkeys’ choice accuracy provided quantitative predictions about subjective confidence. These predictions fit the monkey’s opt-out behavior in our task, indicating that the monkey’s confidence matches the POMDP framework. Prediction of confidence purely based on choice accuracy is a remarkable feat, especially considering systematic discrepancies between the two (Pouget et al., 2016).

Discrepancies between accuracy and confidence have been commonly considered as evidence for suboptimal decision-making or distinct processes that underlie choice and confidence. Our POMDP framework challenges these interpretations by showing that a normative Bayesian decision maker optimizing a reward function elicits the same discrepancies between confidence and accuracy as those identified in humans and experimental animals. We explored five common discrepancies in this paper. Two of them arise from the decision maker’s incomplete knowledge of the environment. The first one is the hard-easy effect, where decision makers are over-confident for difficult choice and under-confident for easy choices (Drugowitsch et al., 2014; Sanders et al., 2016). This effect arises from the decision maker’s uncertainty about the stimulus strength, which results in a regression toward intermediate confidence levels for a Bayesian observer. The second discrepancy is the opposing effects of stimulus volatility on choice and confidence, where subjects become less accurate but paradoxically over-confident about more variable stimuli (Zylberberg et al., 2016). This effect arises from another form of incomplete knowledge about the environment: attribution of the observation noise learned in less volatile environments to newly experienced, more volatile conditions.

The other three discrepancies between accuracy and confidence are explained by our model as arising from the experimenter’s incomplete knowledge of the subjects’ decision-making process. For example, the third discrepancy is based on the observation that confidence reports differ in experiments that interrogate confidence simultaneously with the choice (Kiani et al., 2014) or after the choice (van Den Berg et al., 2016). This difference arises in our model because sequential reports of choice and confidence allow revising one based on information unused for the other. For example, when confidence reports follow the choice, sensory observations in the processing pipeline that were unavailable at the time of the choice could change confidence (van Den Berg et al., 2016).

The fourth discrepancy is the inequality of *d′* and meta-*d′*, which has attracted much attention lately as experimental support for distinct processes underlying choice and confidence (Maniscalco and Lau, 2012). We show that this difference could arise even when a unitary process shapes both choice and confidence, as in our model. A cost-based termination criterion for the decision-making process could affect accuracy and confidence differently. Whereas the overall accuracy decreases due to early termination, confidence can increase especially for incorrect choices, causing unequal *d′* and *d′*. It is therefore impossible to uniquely interpret meta-*d′* in the absence of accurate knowledge about the form of the termination criterion. However, common task designs for measuring choice and confidence often preclude such knowledge.

It is also important to mention that in the presence of observation cost, meta-*d′* depends on the confidence rating threshold in the POMDP model. This sensitivity questions one of the key assumptions in the definition of meta-*d′* — independence of meta-*d′* from the confidence rating criterion — and cautions about interpretations of meta-*d′* results without knowing about the variability of confidence rating thresholds across subjects in an experiment.

The fifth discrepancy that the model explains is the hypothesis that confidence is more strongly influenced by choice-congruent observations than choice-incongruent observations (Zylberberg et al., 2012; Peters et al., 2017). Although these experimental results could indicate post-choice re-weighting of observations for calculation of confidence, they could also arise from the experimenter’s incomplete or inaccurate knowledge of the exact observations used by the decision maker. Many factors could engender such inaccuracy, including neural noise, which is often inaccessible to the experimenter, device noise, which is difficult to eliminate for electrophysiological and imaging techniques, or termination criteria for the decision-making process, which the experimenter may be unaware of or unable to identify.

To clarify our conclusion, we do not imply that dual or hierarchical processes for choice and confidence could not exist. Nor do we exclude the possible existence of mechanisms that revise confidence by post hoc choice-dependent re-weighting of the observations. Rather, we conclude that existing experimental results are insufficient to support such mechanisms as they are also compatible with simpler, more parsimonious mechanisms in which a unitary process underlies both choice and confidence. It is further illuminating that the unitary process explored in this paper is based on POMDPs, a normative Bayesian framework based on expected reward maximization. In light of our POMDP model, existing experimental results should be carefully reconsidered and better experiments should be developed to test the necessity of more complex or disparate mechanisms for choice and confidence.

We applied the POMDP framework to a fixed-duration task where the stimulus duration was controlled by the experimenter. The framework can also be used to model animal and human behaviour in reaction-time tasks Rao (2010). In fact, reaction time tasks might offer better opportunities to study choice and confidence. In fixed-duration tasks, long stimulus durations are least desirable because a multitude of mechanisms, including decision bounds, time-varying attention, or task engagement, could cause partial use of sensory information unbeknownst to the experimenter. Short stimulus durations are not immune to misinterpretations either. Short stimuli can cause neural responses that last longer than the stimulus duration (KovAcs et al., 1995; Kira et al., 2015; Zhou et al., 2018), providing an opportunity for selective sampling. Moreover, short-term mnemonic mechanisms and active revision of choice and confidence provide additional opportunities for dissociating the observations used by the decision-making process from those assumed by the experimenter. Reaction-time task designs where subjects control the stimulus viewing duration, combined with monitoring and manipulation of neural responses in sensory and decision-making circuits, would improve experimental control and enable more accurate interpretation of experimental results (Waskom et al., 2019).

We conclude by noting that simple bounds on decision variables, as employed in traditional models of decision making, might not be sufficient to capture the types of complex policies (mappings of beliefs to actions) required in dynamic environments and in tasks more complex than the random dots task. In such cases, the POMDP model offers a powerful and flexible framework for decision making as it allows (i) arbitrary probability distributions for the prior and observation functions, (ii) arbitrary state transition functions conditioned on the decision maker’s actions, and (iii) policies that are not restricted to bounds on decision variables and that implement arbitrary mappings of beliefs to actions (Rao, 2010; Huang et al., 2012; Khalvati et al., 2019b,a). Testing these more general attributes of the POMDP model in animal and human experiments remains an important direction for future research.

## 4 Methods

### 4.1 Direction discrimination task with post-decision wagering

Complete details of our decision making task are provided in (Kiani and Shadlen, 2009). Briefly, two macaque monkeys were trained to report the net direction of motion of a stimulus of randomly moving dots, a fraction of which moved in a particular direction (Britten et al., 1992). Each trial began with the appearance of a fixation point (FP) on the screen. Shortly after the monkey fixated the FP, two red dots appeared on the two sides of the monitor to indicate the two possible directions of motion in the trial (“direction targets”). After a short delay, the random dots stimulus appeared for 100-900 ms. Motion direction and strength (fraction of coherently moving dots) varied randomly from trial to trial. The random dots stimulus was followed by another delay. Then, on a random half of trials, a third target (called the “sure target”) appeared on the screen in the middle of this delay period. At the end of the delay, the FP disappeared, signaling the monkey to make its choice by making a saccadic eye movement to one of the targets. Choosing the correct direction target (right target for rightward motion and left target for leftward motion) resulted in a large reward (a drop of juice), whereas choosing the incorrect direction target resulted in a timeout. On the trials where the sure target was presented, the monkey could opt out of making a direction discrimination decision by choosing the sure target. The sure target was guaranteed to yield reward but the reward magnitude was smaller than that for the correct direction target (reward ratio, *∼*0.8).

We analyzed the data from the two monkeys separately. Monkeys M1 and M2 contributed 86,622 and 60,733 trials respectively to the dataset.

### 4.2 Model Fits

We used 10 ms time steps in our model fits and simulations because it offered a fine enough temporal resolution to explain the experimental data while keeping the computations manageable. All fits were based on Maximum Likelihood Estimation (MLE).

#### 4.2.1 Model with zero-observation cost

To fit the POMDP model with zero observation cost to the monkey’s accuracy, we found *w*_*z*_ for each monkey’s observation distribution that maximized the likelihood of the monkey’s direction choices on trials without the sure-bet option based on Equation 8. Specifically, choosing the direction *right* with a Bernoulli distribution whose mean is 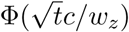 for trials with duration we modeled the probability of *t* and signed coherence *c* (when *c* is negative, the probability of choosing the direction *right* becomes less than .5).

Each monkey’s data were fit separately. For monkey M1, *w*_*z*_ was .90 while for monkey M2, it was 1.69. Based on these *w*_*z*_ values, we estimated the prior belief 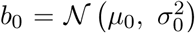 as follows: for any trial with true coherence *c* and duration *t*, we generated a sample from 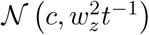; the samples generated from all the trials were used to fit the Gaussian 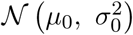 via maximum likelihood estimation (Murphy, 2012).

To calculate *σ*_*z*_, we fit the POMDP model’s belief (Equation 5) to the accuracy in all trials that the sure-bet option was not offered, using *w*_*z*_ and *σ*_0_ estimated as above. In each trial, we calculated 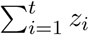, the sum of the observations generated from the actual coherence and the stimulus duration used in that trial. Using the relationship between the sum of observations, *µ*_*t*_ and *σ*_*t*_ in Equation 7 and substituting into Equation 5, we get 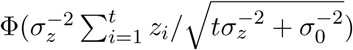 as the subject’s belief about the direction *right*. We calculated a maximum likelihood estimate of *σ*_*z*_ by fitting this belief to the accuracy in all trials where the sure-bet option was not offered. For the fitting, the direction *right* choice was modeled as a Bernoulli distribution whose mean is 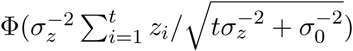, where the *z*_*i*_ were sampled based on the true coherence and duration used in the trials.

The approach described above has the advantage that it can be generalized to other observation distributions and non-zero observation cost. It also suggests the plausibility of the subject learning *σ*_*z*_ without direct access to the actual generative model of observations. For a POMDP with Gaussian prior and observation function and no cost, the average belief for the direction *right* to the true choice can be expressed as a cumulative normal distribution function depending on *t, w*_*z*_, *σ*_0_, and *σ*_*z*_. This means that we could also obtain *σ*_*z*_ without any sampling method.

As explained above, to estimate *σ*_0_, we fit a Gaussian distribution to samples of mean inferred coherence for each trial. This makes the fit independent of *σ*_*z*_. Specifically, if the subject assumes a uniform prior over all coherence values, the belief about the coherence after observing *z*_1_, …, *z*_*t*_ is a Gaussian distribution with a mean 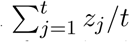 and variance 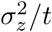 (similar to equation 7 with 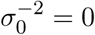). As a result, the distribution of mean inferred coherence is *𝒩* (*tc, tw*_*z*_) for trials with duration *t* and coherence *c*. Therefore, we can estimate the overall distribution of mean inferred coherence, and consequently *σ*_0_ without *σ*_*z*_. One can also try to make the fit more accurate by considering the subject’s uncertainty about the mean coherence, i.e., the variance of 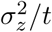 in the belief after observing *z*_1_, …, *z*_*t*_. This, however, requires *σ*_*z*_. To test whether this method improves the fit to data, we started with the value of *σ*_0_ obtained by the fit to mean inferred coherence, estimated *σ*_*z*_ and readjusted *σ*_0_ based on this estimated *σ*_*z*_. The readjusted *σ*_0_ was then used to fit *σ*_*z*_ again. With every such iteration, the change in *σ*_0_ decreased. We repeated this process until the change in *σ*_0_ became less than our precision error. This process converged in less than 5 iterations for both monkeys. However, the readjusted *σ*_0_ values did not improve the goodness of fit of the belief to the monkey’s choice significantly. Nonetheless, we used these more accurate values in our models: *σ*_0_ was .46 and .87, and *σ*_*z*_ was 1.60 and 3.59 for monkey M1 and M2, respectively.

The estimated *σ*_*z*_ was larger than *w*_*z*_ in both monkeys due to the structure of the task. The task involved a discrete set of seven coherence levels, {0%, 1.6%, 3.2%, 6.4%, 12.8%, 25.6%, 51.2%}, and the belief for each direction is the average probability of that direction over these coherence levels (Khalvati et al., 2016; Drugowitsch et al., 2014). The proportion of “low” versus “high” coherence levels in the experiment determines the difference between a POMDP model with continuous coherence values and *σ*_*z*_ = *w*_*z*_, and the true generative model with seven discrete coherence levels. Specifically, in an experiment with a greater number of low coherence trials than high coherence trials, such as ours, the continuous model with *σ*_*z*_ = *w*_*z*_ generates higher confidence compared to a model with the true generative model, resulting in an overall overconfidence for each stimulus duration. Therefore, a continuous model that matches its confidence with a subject’s accuracy for each stimulus duration considers each observation less reliable, i.e., *σ*_*z*_ *> w*_*z*_.

We fit the POMDP model to predict confidence as a function of coherence (motion strength) and duration. To compare the quantitative match of our predictions with the monkey’s post-decision behavior, we used a threshold on confidence that determined whether to choose or reject the sure-bet option when it was available. We fit this threshold by maximizing the likelihood of the monkey’s choices (choosing or rejecting the sure-bet) based on the model’s predicted confidence. The estimated threshold was .63 and .59 for Monkey M1 and Monkey M2, respectively.

Finally, we calculated the model’s prediction of accuracy on the trials that had confidence levels above the threshold. The predicted accuracy was compared to the monkey’s accuracy on trials in which the sure-bet option was available but ignored by the monkey (Fig. 3d). Trials with 0% coherence were removed from this accuracy analysis because a correct direction choice is undefined on those trials and the monkey was rewarded randomly.

#### 4.2.2 Model with non-zero-observation cost

We also tested the POMDP model with non-zero observation cost. This model, with two parameters (*w*_*z*_ and observation cost), was fit to the monkey’s choices on trials without the sure-bet target. Similar to the fitting procedure above for the zero-cost case, *σ*_0_ and *σ*_*z*_ were obtained by an iterative process that fit the average belief to average accuracy for each time step and *σ*_0_ was estimated based on the overall observation distribution. Because there is no closed-form equation for probability of choices in this model (in contrast to the zero-cost case that uses Equation 8), we used grid search for the free parameters and estimated choice probability using particle filtering with 20,000 samples (Thrun et al., 2005). The grid resolution for cost was 10^*−*5^ while for *w*_*z*_, it was 0.01.

### 4.3 One-step Look-Ahead Search as the Optimal Strategy

For our results, we used one-step look-ahead search. Here we show that for an unbiased 2-alternative decision-making task such as ours, one-step look ahead search results in the optimal POMDP policy for a non-decreasing observation cost over time. First, note that due to the symmetry of the task for direction choices, the optimal decision maker picks the choice with the highest belief. This means that when considering whether to terminate or continue acquiring observations, an optimal decision maker compares the observation cost and the resultant expected confidence (belief).

Second, the entropy, i.e. *−b*_*right*_ log(*b*_*right*_)*−b*_*left*_ log(*b*_*left*_), has an inverse relationship with confidence. The expected information gain (i.e., decrease in entropy) decreases with more samples (here observations) (Golovin et al., 2010). As a result, the expected increase in confidence decreases with the number of observations as well. This means that if the expected increase in confidence with one more observation is less than the cost of the observation, the expected increase in confidence with *k* more observations is less than *k* times the cost of one observation. Thus, if the cost of observations is non-decreasing over time, comparing the expected confidence with the cost of an observation at the current time is enough to maximize the expected total reward. In other words, if the next observation is not worth its cost, making more observations would not be worth the cost either. Importantly, this holds for any observation function and state space as long as the probability distribution for observations does not change with time, which is true in our task (coherence does not change within a trial).

### 4.4 Vuong’s Statistical Test

To test whether the monkey’s observation cost was negligible in our task, we used Vuong’s closeness test which compares the goodness of fit of two models, *u*_1_ and *u*_2_, based on their likelihood ratio and number of parameters (Vuong, 1989). With *N* data points in a data set, the *Z*-statistic of this test is:

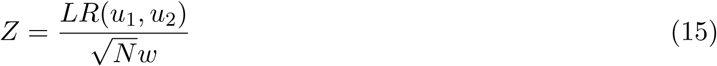

where *LR*(*u*_1_, *u*_2_) = *L*_1_ *− L*_2_ *−* .5(*K*_1_ *− K*_2_) log *N*. *L*_1_ and *L*_2_ are the log likelihoods, *K*_1_ and *K*_2_ are the number of parameters of *u*_1_ and *u*_2_, respectively, and *w* is the mean of the squares of the pointwise log-likelihood-ratios between the two models. We used Vuong’s test to compare the fit of the zero-cost and non-zero-cost POMDP models to our experimental data. There was no significant difference between the two models, even without penalizing the non-zero-cost POMDP model for having one more parameter (i.e., with *LR*(*u*_1_, *u*_2_) = *L*_1_ *− L*_2_).

### 4.5 Simulations for Increased Stimulus Volatility

We used the POMDP model with parameters fit to Monkey M1’s data. For the low volatility regime, the standard deviation of observations was *w*_*z*_ = .9 and the learned standard deviation was *σ*_*z*_ = 1.6. For the high volatility regime, the standard deviation of observations was *w*_*z*_ = 1.5 without changing *σ*_*z*_ or any other parameter in the POMDP model.

### 4.6 Simulations for Simultaneous and Sequential Reports of Choice and Confidence

We used the following POMDP model parameters: *w*_*z*_ = .4, *σ*_*z*_ = .75, and *σ*_0_ = 5 with the 7 discrete coherence levels used in our monkey experiment and a constant observation cost of 2 *×* 10^*−*3^ per 10 ms to simulate the reaction time task with 20,000 trials for each coherence. In the simultaneous report version of the model, both confidence and choice were calculated from the observations received prior to the model reaching its decision termination bounds. In the sequential report version, calculation of confidence continued to be influenced by observations during a 250 ms non-decision time after the choice.

### 4.7 Exploring the Effect of Cost on Sensitivity Measurements and Confidence Report

We compared the POMDP model obtained from Monkey M1 with a model with similar parameters (*w*_*z*_, *σ*_*z*_, and *σ*_0_) but with an observation cost of 10^*−*4^ added to establish decision termination bounds in trials with coherence of 12.8% and duration of 400*ms*. The confidence report was in the form of a binary rating (low or high) with a threshold of 0.63 applied to the belief about the choice. The sensitivity (*d′*) and meta-*d′* were both 1.79 for zero observation cost. Increasing the cost to 10^*−*4^ decreased *d′* to 1.66 and meta-*d′* to 1.23. We used 1 million samples to ensure the results were robust. The code from (Maniscalco and Lau, 2012) was used to calculate meta-*d′*. Qualitatively similar results are obtained for other motion coherence levels and durations.

### 4.8 Prediction Power of Choice-Congruent and Choice-Incongruent Observations

First, we simulated the random dots motion discrimination task with one coherence (*w*_*z*_ = 1, *c* = 10.0%), one duration (800 ms) and a binary confidence rating of low or high with a POMDP model that had an exact model of the world (i.e., with the true *w*_*z*_ and *c*) but used the first *n* observations out of *t* = 80 (step size = 10 ms) observations. For each *n*, the confidence threshold was set to a value that made the probability of high confidence around 0.5.

To generate data points for Figures 6a and 6b, we trained logistic regression classifiers to predict the simulated choices and confidence ratings. 10 million trials were simulated for these analyses to ensure robust and accurate results. Our classifiers were implemented using the `scikit-learn` Python library (Pedregosa et al., 2011). For choice, the features of our classifier were the sum of positive observations and the sum of negative observations throughout each trial, including those beyond the first *n* samples used for simulating choice and confidence. For confidence, the features were the sum of choice-congruent observations and the sum of choice-incongruent observations throughout each trial. For the balanced classifier, to ensure balance of weights, we used a classifier with a feature consisting of the sum of all observations signed according to the choice (positive for choice-congruent and negative for choice-incongruent) as one feature.

For generating Figures S1a and S1b, we repeated the above analysis with the same parameters but with the simulations using all *t* = 80 observations and the classifiers (representing the experimenter) using only *n* of those observations.

For Figures 6c, 6d, S1c, and S1d we added zero mean Gaussian noise with a standard deviation of 1.12 to the observations used in our classifiers to mimic the noisy estimate of a subject’s observations used by an experimenter.

## 5 Acknowledgements

We thank Saleh Esteki, Christina Hatch, Gouki Okazawa, John Sakon, Long Sha, Michael Waskom, and Mike Shadlen for helpful discussions. R.K. acknowledges support from the Simons Collaboration on the Global Brain (542997), National Institutes of Mental Health (R01 MH109180-01), Alfred P. Sloan Foundation, the McKnight Foundation, and a Pew Scholarship in the Biomedical Sciences. R.P.N.R. acknowledges support from the National Institutes of Mental Health (CRCNS 5R01MH112166-03), National Science Foundation (EEC-1028725), Templeton World Charity Foundation, and a CJ and Elizabeth Hwang Endowed Professorship.

