## Supplemental Figure 1 for "Bayesian Inference with Incomplete Knowledge Explains Perceptual Confidence and its Deviations from Accuracy"

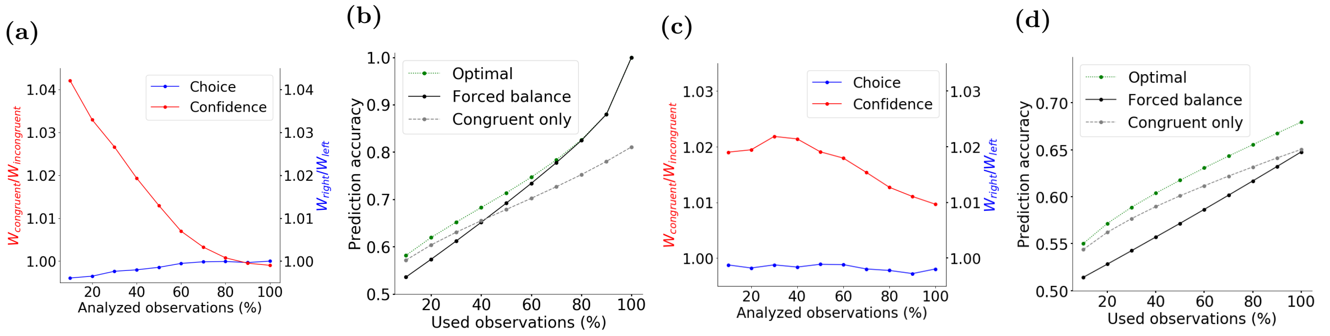


**Figure S1: Sub-sampling the decision maker's observations leads to seemingly higher influence of choice-congruent evidence on confidence ratings.** In section 2.4.5, we demonstrated that when a decision maker uses a subset of available observations unbeknownst to the experimenter, the experimenter is prone to conclude a larger influence of choice-congruent observations on confidence. This artifact arises from misestimation of the congruent and incongruent evidence available to the decision maker at the time of decision. For similar reasons, if the experimenter analyzes only a subset of the observations used by the decision maker, they would artifactually find a larger effect of choice-congruent observations on confidence ratings. Simulation parameters in this figure are similar to those in figure 6, except that the analyzed observations are a fraction of those used by the POMDP model. Note that the POMDP model that simulates the decision maker treats choice-congruent and incongruent observations identically. Therefore, the ground truth is equal weights for those observations, both for choice and confidence ratings. **a)** Choice-congruent observations gain larger weight in a classifier that predicts the decision maker's confidence (low or high) based on the observations analyzed by the experimenter. Note the change of y-axis scale compared to figure 6a, indicating that the weight imbalance here is much smaller than when the decision maker uses less evidence than the experimenter assumes. **b)** Similar to figure 6b. Forcing the classifier to assign identical weights to congruent and incongruent observations reduces classification accuracy. **c-d)** Same as a-b, but with noisy estimates of the decision maker's observations accessible to the experimenter. Such noise reduces the prediction accuracy of the confidence classifier, but more important, it also causes imbalanced weights in the optimal classifier that persist even when the classifier is based on all the observations used by the decision maker. Further, the contrast between the optimal classifier and the classifier with balanced weights increases compared to panel a. The noise comes from a zero-mean Gaussian distribution with a variance 25% larger than $w_z^2$, similar to figures 6c and 6d.
